# Gaps and Advances in Long-Term Monitoring of Antarctic Near-Shore and Terrestrial Ecosystems

**DOI:** 10.1101/2025.04.03.647064

**Authors:** Abigail Borgmeier, Dana Bergstrom, S. Craig Cary, Peter Convey, Vonda Cummings, Dolores Deregibus, Soon Gyu Hong, Charles K. Lee, Hyoungseok Lee, Marcela Libertelli, Sharon Robinson, Stefano Schiaparelli, Emmanuelle Sultan, Megumu Tsujimoto, Elie Verleyen, Byron J. Adams

## Abstract

Environmental change due to greenhouse gas emissions is affecting ecosystems globally; in the polar regions in particular there is already significant evidence of change. The Antarctic Near-Shore and Terrestrial Observation System (ANTOS) aims to establish a cross-continent, cross-national observing network to assess environmental variability and change in the southern polar region. To understand how near-shore and terrestrial Antarctic ecosystems have and will continue to be impacted by anthropogenic environmental changes, a comprehensive review of current long-term monitoring efforts, and two surveys targeting Antarctic researchers, were carried out to evaluate existing monitoring efforts, environmental changes recorded, and to identify areas currently lacking sufficient observations. Results indicate that most data collection is manual and intra-annual, with significant long-term monitoring concentrated in regions with already established infrastructure. The surveys highlight the urgent need for comprehensive coverage of the Antarctic’s rapidly changing ecosystems using standardized monitoring protocols and increased collaboration. We recommend prioritizing areas experiencing rapid climatic changes and leveraging existing infrastructure to minimize environmental impact of monitoring activities and enhance data comparability across sites.

## Introduction

Environmental change due to greenhouse gas emissions is affecting ecosystems globally (Hoegh Guldberg et al., 2018; Keeling, 2008). Long-term monitoring of various environments has revealed ecosystem impacts after years of climate change, which include species’ range shifts, local and regional extinctions, the introduction of non-indigenous species, homogenization of biota across bioregions, and other biotic changes (Gossner et al., 2023; Morley et al., 2020; Xian et al., 2023). The numerous effects due to climate change have rapidly increased in frequency and intensity in the last several decades (Lee et al., 2023), and now include the discussion of tipping points (Clark et al., 2013; Heinze et al., 2021; Lenton et al., 2019; Muthukrishnan et al., 2022), as environments change dramatically, unpredictably, and perhaps irrevocably. Many insights into environmental changes driven by climate change can only be discovered through research based on long-term monitoring (Bauman et al., 2022; Oliver et al., 2015; Rull, 2014). Observations from some long-term monitoring studies have led to changes in management practices, which have been crucial to the maintenance of natural spaces following climate fluctuations (Brienen et al., 2015; Smith et al., 2016).

### Importance of long-term monitoring

The Antarctic region is one of the most sensitive locations to warming on the planet (Mulvaney et al., 2012) and has been used as a barometer for the world to measure how climate change has and will impact delicate natural ecosystems (Chown et al., 2022). In the Antarctic, the goal of long-term monitoring is not only to better manage land preservation, but also to examine how ecosystems adapt and change due to climate change (Cimino et al., 2023; Piazza et al., 2020). Long-term monitoring has already revealed several potential physical drivers of ecological change in Antarctic ecosystems. Stream gauges in the Dry Valleys region of Victoria Land, Antarctica documented significant changes in hydrological connectivity and found that the majority of water in streams comes from glacier melt, which is expected to increase in the coming years (Gooseff et al., 2022). Four decades of monitoring of the Antarctic ice-sheet showed that the rate of ice mass loss is increasing over time (Rignot et al., 2019).

The goals of these projects could not have been met without an established baseline of the biotic and abiotic factors governing ecosystems. Without such a baseline, unusual weather events might be misinterpreted as isolated anomalies rather than components of larger patterns characterizing the continent. Antarctica experiences both ‘pulse’ events (acute and discrete events), and ‘press’ events (that occur continuously through time) (Inamine et al., 2022). Continuous monitoring captures both types, whereas infrequent or sporadic observations generally detect only press events. Therefore, long-term monitoring in Antarctica is essential for documenting the environmental impact of climatic changes on the continent. This endeavor requires collaboration across the national scientific programs working in Antarctica to ensure that monitoring data are comparable and usable by all scientists working on and around the continent.

### Criteria for establishing long-term monitoring in Antarctica

Antarctic ecosystems have changed substantially over the last several decades, largely due to anthropogenically driven climate change (Bornman et al., 2019; Chown et al., 2022; Grant et al., 2021; Lee et al., 2017; Stark et al., 2014; Turner et al., 2014). Identifying the most critical areas for monitoring and pinpointing the gaps in current efforts has become paramount.

There are several challenges specific to ecological monitoring in Antarctica, but also unique opportunities. One of the opportunities is that the continent has been lauded for scientific research as a “natural laboratory” due to the pristine environment which has been untouched by humans until recent decades (Laybourn-Parry & Pearce, 2007). Therefore, Antarctic ecosystems are uniquely sensitive to disturbance and change due to human presence, including research presence, making it challenging to establish monitoring programs (Barnes & Conlan, 2007; Kiernan & McConnell, 2001). Additionally, it is important to consider the price, difficulty, and resources necessary, and the potential environmental impacts, when implementing and maintaining long-term monitoring projects in such remote and inaccessible areas, and particularly in locations which are not currently being supported for research.

### The Antarctic Near-Shore and Terrestrial Observation System

The Antarctic Near-Shore and Terrestrial Observation System (ANTOS) is a SCAR expert-group whose goal is to “coordinate a cross-continent and cross-national program-scale assessment of environmental variability and change” (Cary & Cummings, 2022; Piazza et al., 2019). To achieve this goal, ANTOS is working to identify nearshore marine and terrestrial locations on the Antarctic continent which are at high risk of change in the coming decades, where there is interest from the scientific community, or where there are no current monitoring programs in place. ANTOS aims to organize the formation of a regulated and wide-reaching network of long-term monitoring sites on the Antarctic continent and its surrounding coastal environment. The group conducted two surveys targeting Antarctic researchers from a variety of different scientific programs, to assess current monitoring efforts and the environmental changes documented. In this manuscript we describe the surveys conducted, identify gaps in current long-term monitoring initiatives on/around the Antarctic continent, and recommend the most important locations for implementing new monitoring efforts.

## Methods

### Survey methodology and data collection

ANTOS conducted two surveys to capture current monitoring efforts and identify gaps in near-shore and terrestrial Antarctic ecosystem monitoring.

#### Survey I

Survey I was designed to canvas a wide variety of respondents to ensure a representative overview of biological data collection on and around the continent. The full list of questions is included in Supplementary Table 1. Respondents were asked to give their name, identify the national program which supports their research, their research discipline, and where they collect data. They were asked to provide information on collection, duration and frequency of data collection. Respondents were also asked to suggest priority locations for monitoring, which were compiled into a list of areas of interest for ongoing and/or future monitoring. The survey was sent to the Antarctic life sciences database, which is an opt-in mailing list organized by the SCAR Life Sciences Group (https://scar.org/science/life/lifescience). A total of 388 responses were recorded. Of the total responses, 111 individuals responded in completion and were included in the survey analysis.

**Table 1.**
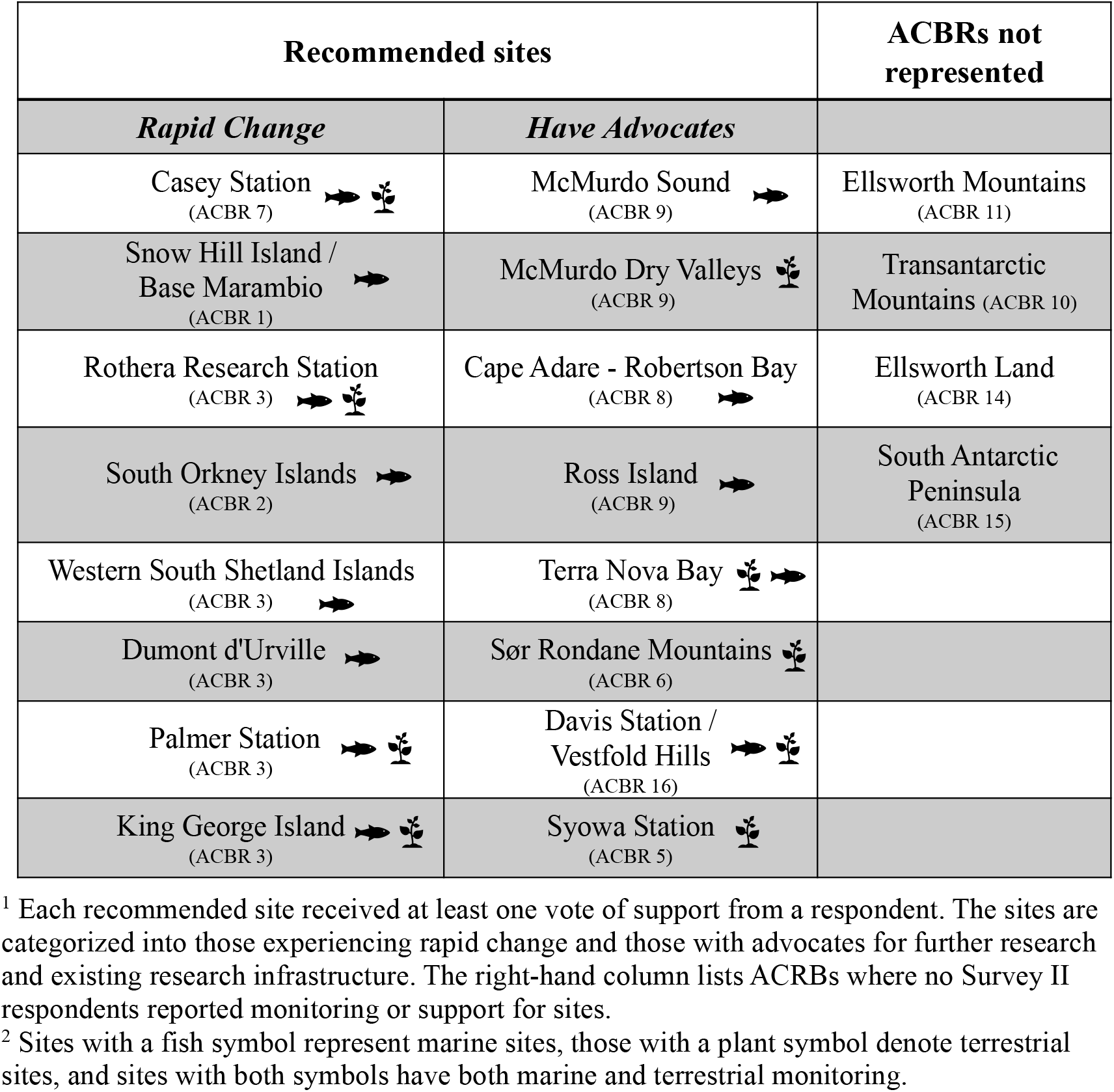
Table of recommended sites based on Survey II respondents.^1,2^.

#### Survey II

The primary goal of Survey II was to assess which sites on the areas of interest list, generated from the responses in Survey I, have the greatest demand for monitoring, and which of the sites are most vulnerable to change or are unique from other sites on the continent. All questions in Survey II are listed in Supplementary Table 2. The respondents were asked to give GPS coordinates of all their current monitoring sites, and describe the type, duration, and frequency of data collection. They were asked to identify the environmental and ecological factors driving changes in their research sites. Additionally, respondents were asked to prioritize the areas of interest for ANTOS monitoring by giving a vote for one or more site on the areas of interest list generated in Survey I. The respondents were asked to defend that vote by identifying what physical or biological attributes are most unique to that site, and why those attributes are valuable to the overarching ANTOS purpose of monitoring key variables that drive ecosystem change. The survey was sent to all respondents from Survey I. Thirty-two individuals from various biological sciences and national research programs responded to Survey II.

### Using existing bioregionalization frameworks

To interpret the survey results, and help visualize monitoring and knowledge gaps, we used the Antarctic Conservation Biogeographic Regions, which was redesigned and updated by Terauds & Lee, 2016). The Antarctic Conservation Biogeographic Regions (ACBRs) divide the Antarctic continent into 16 biologically distinct units (Terauds et al., 2012). These units have been used in the past to describe Antarctic biodiversity ranges (Chown et al., 2015), as a method of evaluating conservation efforts of Antarctic biodiversity (Hughes & Grant, 2018; Hughes et al., 2016), and to illustrate consequences to biodiversity following anthropogenic disruption (Hughes et al., 2019). While there are recognized limitations in the use of ACBRs; i.e. (i) they cannot perfectly describe the biological complexity on the continent, (ii) climate change and human-mediated dispersal is predicted to increase homogenization across bioregions as the bioregions become less differentiated from neighboring regions (Convey & Peck, 2019; Hughes et al., 2019), and (iii) the ACBRs were designed using primarily a limited range of non-microbial terrestrial taxa; near-shore and microbial diversity is underrepresented (Varliero et al., 2024), they are nevertheless a useful strategic framework for selecting new long-term monitoring sites.

Multiple Marine Protected Areas (MPAs) have been established around Antarctica for management and conservation (Hawkey et al., 2013; Brooks et al. 2020), although this process has progressed slowly with much geopolitical obstruction within CCAMLR (Brooks, 2013). Ecoregions have been defined within some but not all of the MPAs (Brooks et al., 2020; Koubbi et al., 2016; Koubbi et al., 2010), and therefore have not been used in our long-term monitoring site selection process.

## Results

### Survey I

Of the 111 complete responses to Survey I, 21 countries were represented, with 80% of those responses being from biological science researchers, 8% from physical sciences, and 5% from geological sciences. Twenty-eight sites were included on the areas of interest list for potential future monitoring sites (Figure 1). These sites represent locations where both terrestrial and near-shore marine ecosystems could be monitored. Forty-nine percent of data is being collected periodically through instrumentation. A suite of abiotic and biotic factors are being measured. The most common abiotic factors measured are air temperature (80% of respondents) and water temperature (47% of respondents). The most common biotic factors being measured are biodiversity (65% of respondents) and flora/fauna (49% of respondents). Of any factor being measured, 34% of respondents reported that their site has been monitored in some capacity for 21-50 years, and 32% reported monitoring occurring for 11-20 years.

**Figure 1.**
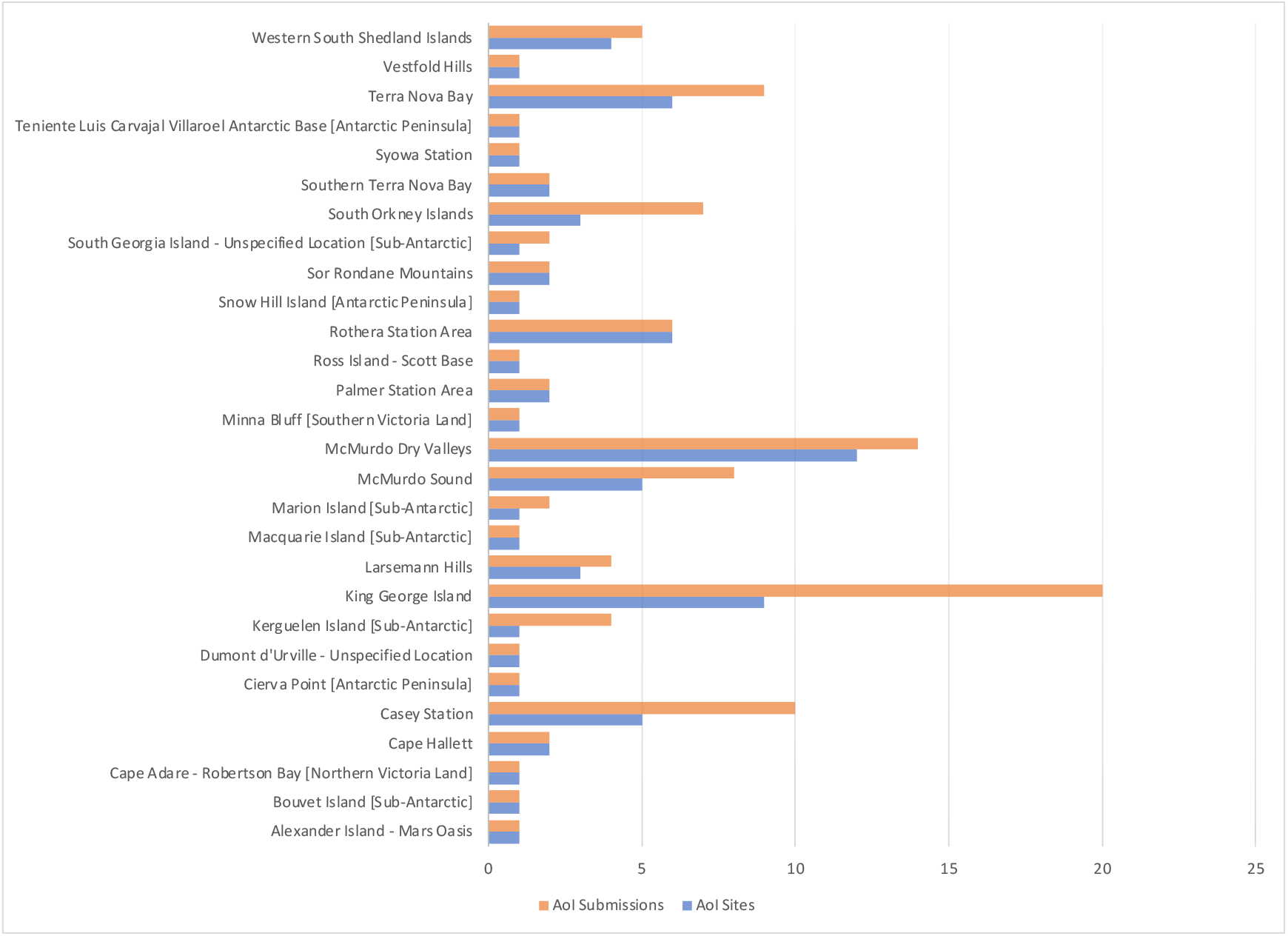
Areas of interest (AoI) compiled from responses to Survey I. For each region listed on the y-axis, the number of sites proposed is the length of the blue bar. The number of votes for any site within that region is the length of the orange bar.

**Figure 2.**
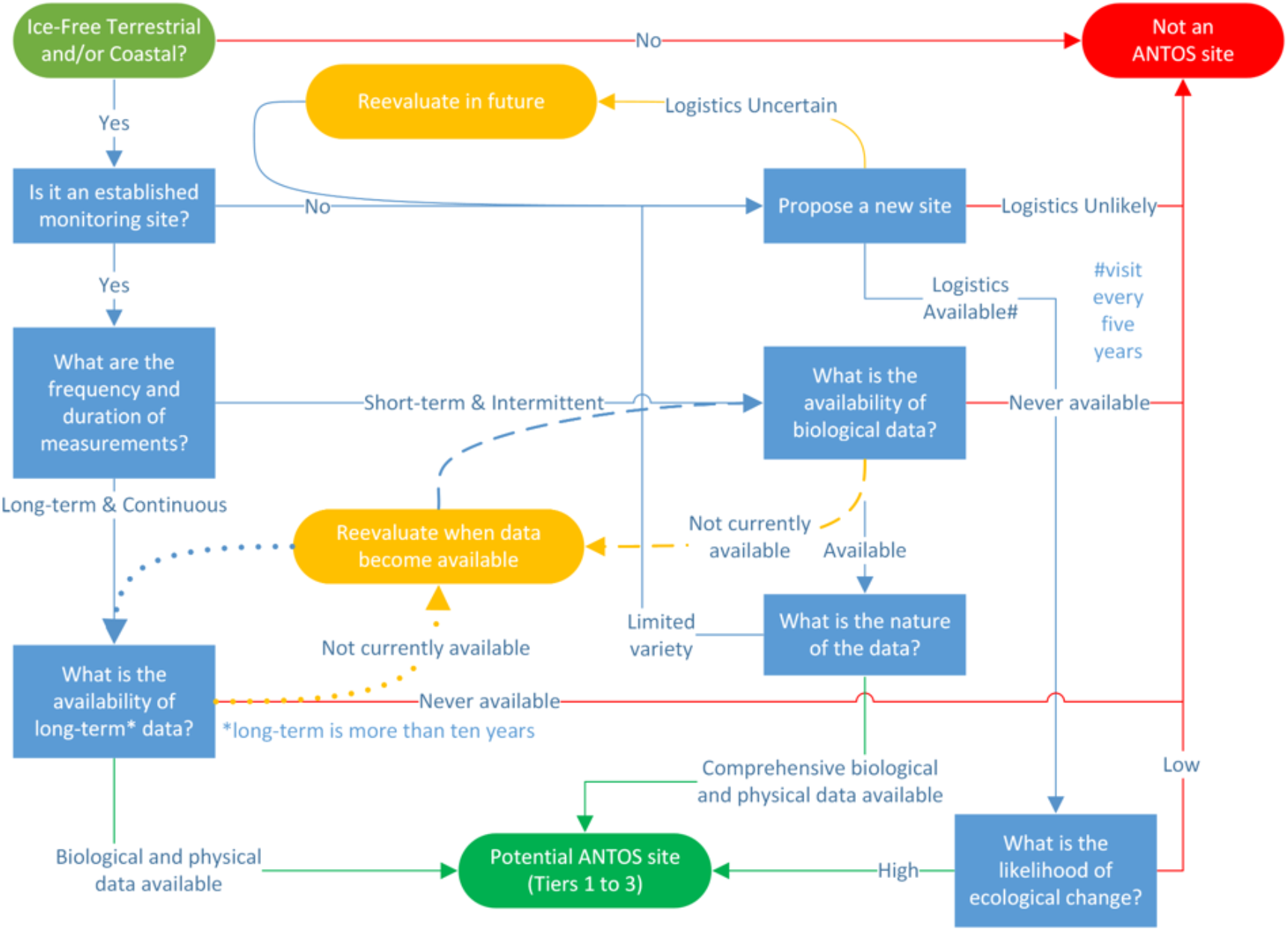
Decision-making framework for ANTOS candidate site selection. The decision tree provides a structured approach to evaluating potential Antarctic Near-Shore and Terrestrial Observation System (ANTOS) sites. The process prioritizes locations based on three criteria: ecological significance, evidence of rapid environmental change, and logistical feasibility for sustained monitoring.

### Survey II

Based on the results of Survey II, most of the data currently being collected is collected manually (versus automatically) and on annual or shorter timescales (versus continuously). There are a variety of both biotic and abiotic factors being measured across the continent (Figure 3). Wind speed and direction have been measured in some locations for the longest period of time, with one respondent reporting that this factor has been measured at their site for over 50 years. Thirteen out of the 34 factors surveyed have been collected somewhere on the Antarctic continent for 21-50 years. In all monitored locations, a mix of biotic and abiotic factors are being measured.

**Figure 3.**
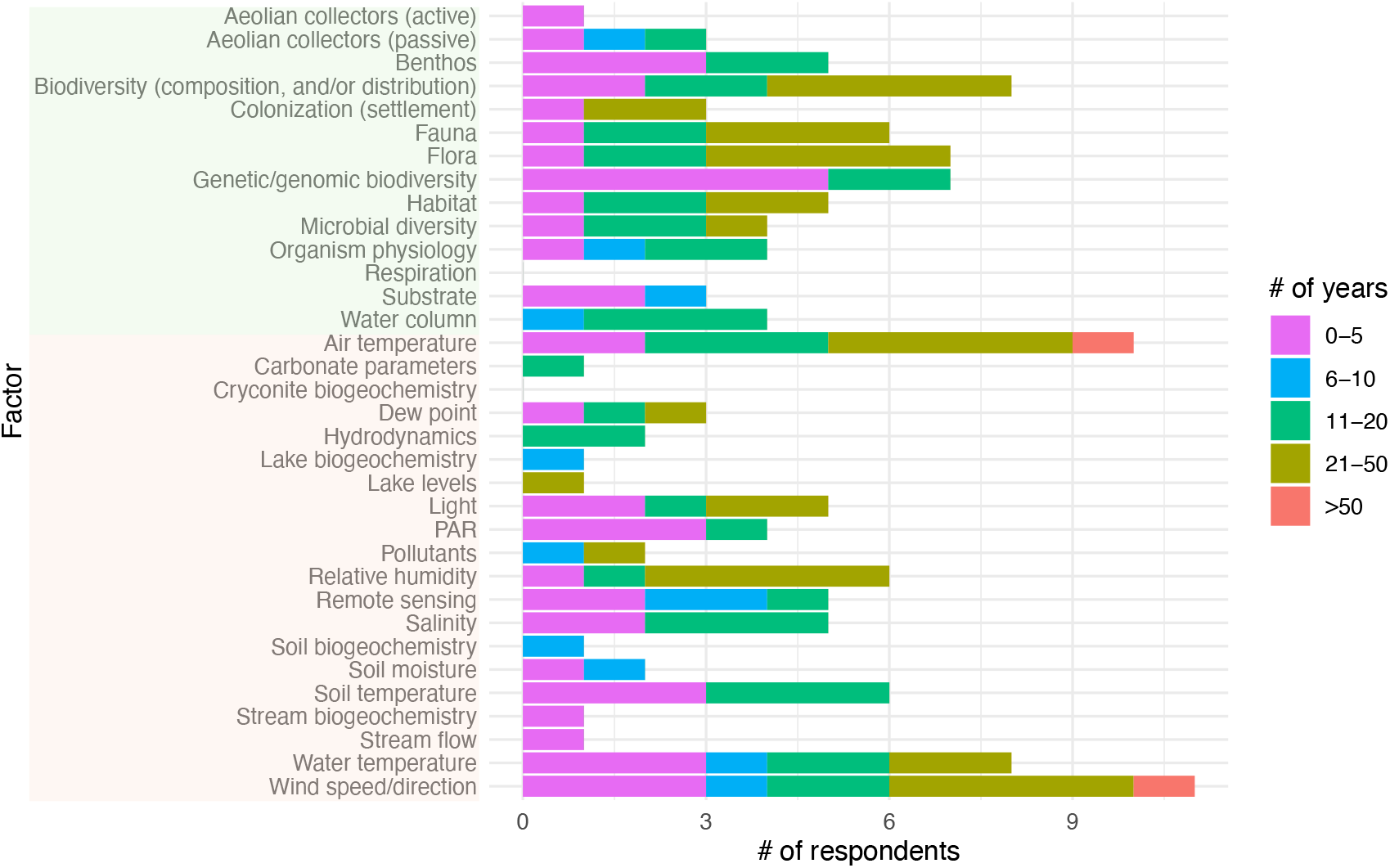
Abiotic and biotic factors monitored across the Antarctic, and the length of time they have been monitored, compiled from results of Survey II. Bar lengths represent the number of respondents who reported monitoring each factor for the indicated number of years. Abiotic factors are in red font, biotic factors are in green font.

Biotic and abiotic factors are being monitored from most, but not all ACBR regions in Antarctica (Figure 4). Sites on the Antarctic Peninsula and Scotia Arc archipelagoes, southern Victoria Land, and Casey Station in East Antarctica have been monitored for over fifty years, making these sites the longest monitored in Antarctica. All Marine Protected Areas are represented by monitoring reported by at least one respondent to Survey II. The locations where monitoring has taken place in the past largely represent active research stations and their surrounding areas. Thirteen out of the 16 ACBRs contain at least one active research station or permanent infrastructure (Terauds & Lee, 2016).

**Figure 4.**
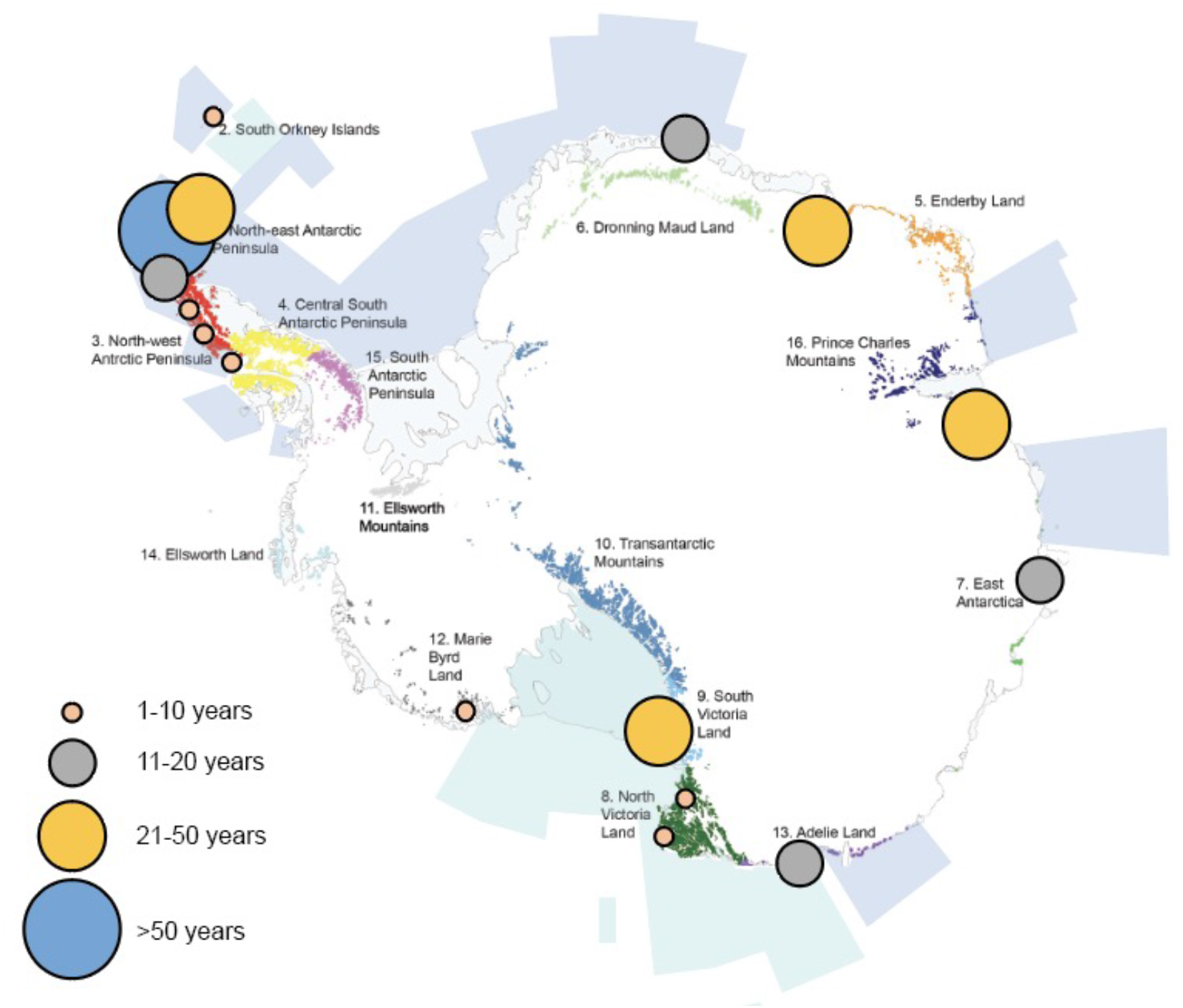
Map of Antarctic Conservation Biogeographic Areas (ACBRs), denoting areas of unique ecological significance (Terauds & Lee 2016). Proposed Marine Protected Areas (MPAs) are shown in dark blue (C. M. Brooks et al., 2020) and current Marine Protected Areas are shown in light blue. Current monitoring efforts are indicated by dots, with the size of each dot indicating the number of years monitoring has occurred in that area. Map modified from on Terrauds & Lee (2016).

The sites on the areas of interest list, generated by Survey I, which did or did not receive a vote of support in Survey II are listed in Table 1. The recommended sites, which had at least one advocate from the Antarctic research community for an ANTOS monitoring node to be built at that site, represent a mix of terrestrial and near-shore marine monitoring sites, and cover most of the ACBRs on the continent. The list of recommended sites are further categorized into sites which are currently experiencing rapid climatic change, and sites which have existing infrastructure or a history or long-term monitoring already underway at that site (Table 1). The ACBRs which were not represented by a vote of support in Survey II are also highlighted (Table 1).

### ANTOS decision tree

After candidate locations have been identified for future ANTOS monitoring sites, the ANTOS decision tree can be utilized to determine if each candidate site is appropriate for the implementation of an ANTOS monitoring node (Figure 2). The decision tree walks through all considerations necessary for determining if a proposed monitoring location falls within the goals of the ANTOS network. Key steps guide users from site identification to final evaluation, supporting a standardized selection across diverse Antarctic regions to ensure effective and collaborative long-term monitoring.

## Discussion

We proposed three criteria which are helpful in determining which sites in Antarctica are appropriate for continued or new monitoring programs. These criteria include 1. evidence of rapid climatic change in the ecosystem, 2. persistent interest in studying the location, and 3. infrastructure present to support long-term monitoring of the site. These criteria are designed to help identify significant deficiencies in long-term monitoring on the Antarctic continent and to determine the most important locations for implementing new monitoring efforts. The proposed network of monitoring sites has the potential to increase cross-site comparisons (Magnuson & Waide, 2021) and reduce the overall human footprint in fragile ecosystems by building a comprehensive data collection and human network. Implementing these monitoring sites will follow ANTOS guidelines, ensuring that data collection is standardized, making the findings directly and easily comparable across all locations (Kao et al., 2012).

### Recommended sites for long-term monitoring

#### Sites experiencing rapid change

Most of the sites which are currently experiencing rapid climactic changes are located in the Antarctic Peninsula region. The Antarctic Peninsula is undergoing, and predicted to continue to experience, rapid warming and loss of ice-covered land (Cannone et al., 2022; Convey et al., 2009; Lee et al., 2017; Turner et al., 2014; Vaughan et al., 2003), glacial retreat and reductions in sea ice cover (Griffiths et al., in press; Gutt et al., 2021). In addition to climate change, the Peninsula is also threatened by considerable densities of research stations and research activity, increased tourism, and introduction of non-indigenous species (Convey & Peck, 2019; Siegert et al., 2019; Tejedo et al., 2022). Rothera Research Station, South Orkney Islands, Western South Shetland Islands, Palmer Station, King George Island/Isla 25 de Mayo, and Snow Hill Island are all recommended sites which are undergoing rapid climatic change. For example, off of the southern Antarctica Peninsula, rising temperatures are predicted to significantly increase fungal diversity in the coming decades (Newsham et al., 2016).

One of the primary terrestrial changes in the Antarctic Peninsula is the increasing amount of newly ice-free land. It has been predicted that, under a current ‘business as usual’ climate warming scenario, over 17,267 km^2^ of new ice-free area will emerge by the end of the century across the continent, with the majority of this increase (a 300% increase regionally) being located in the Peninsula (Lee et al., 2017). The emergence of new ice-free land is a highly significant ecosystem change, allowing for expansion of vegetation, for new species to colonize the area (Cannone et al., 2022), increasing connectivity between previously isolated areas, and increased threat of introduced species (Lee, Waterman, et al., 2022). The near-shore marine ecosystems on the Antarctic Peninsula have also experienced rising ocean temperatures, ice-shelf collapse, and species diversity decline (Ingels et al., 2021; Rogers et al., 2020). Marine species’ ranges are predicted to shift in response to warmer ocean temperatures and reduced levels of sea ice (Barnes et al., 2024; Barnes & Peck, 2008; Swadling et al., 2023), and especially, in coastal areas significant shifts in an Antarctic benthic communities have already been reported and been linked to ongoing climate change (Clark et al., 2013; Deregibus et al., 2023; Quartino et al., 2013).

While most of the Antarctic continent is expected to experience less extreme warming compared to the Peninsula, regions in East Antarctica, including Casey Station, have experienced recent heatwave extreme events (Robinson et al., 2020). Much of East Antarctica has avoided to date the surface air temperature increases that the rest of the globe is experiencing. This is likely a linked consequence of ozone depletion, but that effect will likely reduce in the future (Robinson & Erickson III, 2015; Robinson et al., 2020). Even so, the East Antarctic environment is changing and documented change in biodiversity is occurring, with species turnover and decreased health observed in plant communities (Bergstrom et al., 2021; Robinson et al., 2018). Increasingly frequent extreme temperature events will affect the terrestrial and marine biodiversity of the continental Antarctic region (Barrett et al., 2024; Bergstrom et al., 2021; Robinson et al., 2020; Wille et al., 2024).

Rapid changes in the Antarctic Peninsula and in continental Antarctica demand for rapid action in establishing long-term ecosystem monitoring. Distinguishing between intra- and inter-annual variation and climate, and establishing a baseline of ecosystem fluctuations is essential to capturing the unusual and extreme fluctuations that these regions are already experiencing. Standardized and consistent monitoring is necessary to capture the current abiotic and biotic factors affected by rapid warming and extreme ecosystem change.

#### Sites with existing infrastructure and support

The list of recommended sites from Survey II includes sites with a large amount of existing research infrastructure and strong advocates for continued scientific research and ecosystem monitoring (Table 1). Establishing new monitoring sites in locations with existing infrastructure is important because known ecosystem disturbance in Antarctica results in large part from national scientific research endeavors (Coetzee & Chown, 2016; Stark et al., 2014). Long-term monitoring of various aspects of the ecosystem have been conducted in the McMurdo Dry Valleys, McMurdo Sound, and Cape Hallett for nearly 30 years (Gooseff et al., 2022). Similarly, the Sør Rondane Mountains and Davis Station/Vestfold Hills have existing research stations and active scientific research (Savaglia et al., 2024). Establishing ANTOS monitoring nodes at these sites would be more feasible compared to remote locations because they are already frequently visited, resulting in limited additional disturbance to ecosystems. While monitoring has occurred in these locations for many years, addition of ANTOS systems would allow for data collected on biotic and abiotic factors to be directly comparable to that from other locations.

#### Gaps in current long-term monitoring efforts

Based on the responses to Survey II, there are several ACBRs not represented by existing long-term monitoring and that received a vote in support of establishing an ANTOS long-term monitoring node. These are largely in remote locations on the Antarctic continent where there are not frequent or active research projects, and where there is little existing research infrastructure. However, these locations are still under threat of experiencing ecosystem change, and therefore represent gaps in our current long-term monitoring efforts (Lee, Terauds, et al., 2022; Liu et al., 2022). The ACBR regions which are not represented by current monitoring efforts include the Ellsworth Mountains, the Transantarctic Mountains, Ellsworth Land, South Antarctic Peninsula, and Enderby Land (Table 1). However, given the geographic scope and remote nature of research and monitoring in Antarctica, it may be difficult to support biodiversity surveys or maintenance of remote monitoring systems in these locations.

Some of the ACBRs which are not currently being monitored are already experiencing climatic change in very similar ways to other locations on the Antarctic continent which have been consistently monitored for decades (Griffiths et al., 2017; Lee et al., 2017). For instance, predictions indicate that Ellsworth Land will see a similar increase in the amount of new ice-free land as the Peninsula (Lee et al., 2017). There are several locations along the Antarctic Peninsula which are experiencing ice-shelf collapse and glacial retreat (Morley et al., 2020; Stammerjohn et al., 2012), resulting in shifts in the marine benthic communities (Sahade et al., 2015). Ice-shelf loss is predicted to continue to increase in the Antarctic Peninsula, potentially affecting Ellsworth Land and other locations (Kennicutt et al., 2019). Despite the logistical challenges, the loss of terrestrial ice and ice-shelves makes Ellsworth Land a high-priority location for establishing new long-term monitoring sites. This need is amplified due to the lack of existing infrastructure and support from one or more national research programs.

## Conclusions

ANTOS is a proposed network of standardized monitoring nodes, which will establish a baseline of ecosystem functioning and capture variability and change (including shifts) in a suite of biotic and abiotic factors due to anthropogenic disturbance and climate change. This network of long-term monitoring is designed to allow all measurements to be directly comparable between locations, encouraging collaboration and cross-site comparisons across and around the continent. The implementation of this network will fill major gaps within current monitoring efforts and ensure that the full breadth of terrestrial and marine ecosystem functioning in Antarctica is captured within it. Building and maintaining new monitoring nodes will require commitment of and collaboration between national research programs, existing research stations, and individual investigators. While long term ecological observations are challenging to fund (Malhi et al., 2020), they are essential to understanding, managing, and mitigating the impacts of global change to these unique Antarctic ecosystems. It has been well documented that there are already climate extremes occurring on the continent and in the surrounding ocean, which necessitates urgent action to capture the full consequences of climate change on the vast and unique diversity of Antarctic life.

## Notes

### Competing Interest Statement

The authors have declared no competing interest.

